# Unbiased whole genome comparison of *Pan paniscus* (bonobo) and *Homo sapiens* (human) through a novel sequence match-based approach

**DOI:** 10.1101/2025.06.30.659717

**Authors:** Christof Rosler, Derick DuFriend, Evonn Annor, Rita Njoroge, Elena Davis, Ywahae Law Anthem, Celestino Velásquez, William P. Ranahan, Matthew H. Goelzer, Stephen Wheat, Julianna A. Goelzer

## Abstract

Due to technical and computational limitations, original attempts to compare humans to other non-human primates (NHP) were restricted to specific gene and protein comparisons. With the advances in supercomputing and whole genome sequencing technology, these studies can be revisited to explore entire genomes unbiasedly. A novel alignment-dependent homology algorithm that utilizes a linear search-based approach to find segments of homolog sequences based on a given length of word size, ranging from 32 bp to 1000 bp, was used to perform whole genome comparisons. These sequences were then compared over each chromosome of both the target and control species. Chromosome similarities between *Pan paniscus* and *Homo sapiens* varied greatly across various chromosomes. At 32-bp granularity, chromosome 3 showed the highest similarity (91.96%), while chromosome 5 showed the lowest similarity (59.66%). Overall, this indicates that while there are significant similarities in the anatomical structures, physiological structures, and protein similarities, there are significant differences in the genomic code between the two species. Additionally, not all sequences are conserved equally, underscoring the need to study the role that gene duplications, transpositions, and horizontal gene transfer may play in species divergence.

## Introduction

Preliminary attempts at understanding homology between species utilized anatomical, physiological, and protein similarities. With the sequencing of the human genome and subsequent animal genomes, comparative biology has begun to rely heavily on genomic homology to determine relatedness. Early attempts via DNA-DNA hybridization utilizing genomic sequences for homology showed approximately 1.5% genomic divergence between humans and bonobos (Sibley et al., 1990). However, these early studies focused on hand-selected shared genes between species and singular chromosomes, such as human chromosome 21 and chimpanzee chromosome 22 (Osada & Wu, 2005). These studies identify significant homology between the target genes with only minor mutational differences between human and non-human primates (NHP). While these studies reveal similarities between humans and NHP, the limited scope of the homology investigation, being that of singular genes and singular chromosomes, excludes large regions of the genome from the comparisons. Large portions of higher-order mammalian genomes contain non-protein-coding regions that contain sequences vital for gene regulation, such as transcription start sites (TSS), CpG islands, and trans-element binding sites. Additionally, when examining 44 specific homologous proteins shared by humans and chimpanzees, prior studies revealed substantial similarity in their amino acid sequences, leading to an overall protein similarity estimation of 99% (King & Wilson, 1975). This finding contributed to the widely accepted notion of a close evolutionary relationship between the two species. However, this study also revealed that 50% of the proteins analyzed exhibited similarity between species, while the remaining 50% displayed differences. Therefore, the estimation of genetic similarities between species can vary depending on the specific genes or proteins under examination. As genetic similarity can be assessed at multiple levels, it is evident that similarities in amino acid sequences do not directly reflect the overall genetic similarity at the nucleotide sequence level.

In 1984, Charles Sibley and Jon Ahlquist set out to address the uncertainty in phylogenetic branching of the Hominidae family. Their first conclusion was a significant alteration to the phylogenetic tree. *Gorilla gorilla* diverged first, placing *Homo sapiens* and *Pan paniscus* together, breaking the previous belief of a trichotomy divergence (Sibley & Ahlquist, 1984). Their second major claim came in their reanalysis of the data, where they conducted a total of 514 DNA-DNA hybridizations, producing multiple possible matrixes of ΔT_50_H melting curves. Upon removing poor melting curve results due to temperature overshoots or excessive short tracer fragments, they again supported their original divergence claim (Sibley et. al., 1990; Williams & Goodman, 1989). The main discovery of this work stated that the percent divergence between *H. sapiens* and *P. paniscus* is 1.6%. However, this number is not what is routinely referenced in phylogenetic articles. Rather, the assumed non-divergent 98.4% is often quoted and has become a paradigm for phylogenetic and homologic studies in the past 30 years (Sibley et al., 1990).

Comparative genomic algorithms utilized in determining homology are classified into two main groups: alignment-independent and alignment-dependent. Nucleotide sequence comparisons require the use of computer algorithms to identify homologous sequences. A reference genome, such as *H. sapiens* GRCh38.p13 or *Mus musculus* GRCm38, is used to compare a specified sequence. However, how the reference genome is utilized in the homology search is different between alignment-dependent and alignment-independent algorithms. Alignment-dependent algorithms align a specified sequence to the reference genome directly, where the nucleotide sequences are linearly matched, base pair (bp) to base pair. Any matched base pairs are conserved, and unmatched base pairs are non-conserved. The main advantage of alignment-dependent algorithms is that they can produce mapped genome sequences, providing insight into the conserved genomic structure of the compared genomes and helping to identify single-point mutations that can have major effects on gene regulation and protein synthesis. Notably, a study examining 348 genes revealed that those classified as transcription factors exhibited 47% more amino acid changes in human genes compared to their chimpanzee orthologues, indicating faster divergences in proteins associated with regulating gene expression (Mikkelsen, 2005). This finding emphasizes the significant role of regulatory variations in shaping protein expression and function, underscoring the complexity of proper genome regulation and protein synthesis and highlighting that DNA is not always conserved equally. Performing alignment-dependent search algorithms is computationally taxing and time-consuming, especially when applied to whole genomes. On the other hand, in alignment-independent algorithms, a sequence of nucleotides is not directly matched to a homologous sequence in a reference genome. Instead, alignment-independent algorithms determine homology without producing any alignment. This has the distinct advantage of requiring less computational power and time to compute, with some algorithms only requiring O(n) search time (Haubold, 2014). Alignment-independent searches also do not require the assumption that both compared sequences have the same linear length, allowing for the analysis of different-length sequences. Unfortunately, as per its name, alignment-independent algorithms do not provide any information related to genome mapping, diminishing its effectiveness in revealing informative data relevant to genomic trees.

With advancements in access to supercomputers, alignment-dependent homology algorithms can be more accessible for research purposes. Supercomputers also allow for the creation of homology alignment algorithms that would normally be technologically taxing and time-consuming on a personal computer. Realizing these advantages, an alignment-dependent homology algorithm was developed by utilizing the ORU Research Computing and Analytics (ORCA) TITAN supercomputer. The alignment-dependent homology algorithm, GeneCompare, utilizes a linear search-based approach to find segments of homolog sequences based on a given length of word size, ranging from 32 bp to 2000 bp, that were compared over each chromosome of both the target species and control species. This novel approach to genomic homology aims to reveal true base-pair-to-base-pair homology between two species without bias towards chromosome length or protein-coding sequences.

## Materials and Methods

### GeneCompare

The Oral Roberts University (ORU) GeneCompare program performs literal pattern matching of two FASTA files of any length. It finds matches of strings of a minimum length, *m,* regardless of where the matching string is found in the two files. For a given match, the position of the match is recorded, along with the length of the match. It outputs two binary files of 8-bit integers, one integer for each character of the corresponding input file. The integers are 0 for “no match” and 1 for “match.”

To perform a whole-genome comparison, GeneCompare is run for each chromosome for the “base” genome against each chromosome of the “target” genome, resulting in 2*bt* files, where *b* is the number of base chromosomes, and *t* is the number of target chromosomes. Scripts are developed to cause each pairwise comparison to be submitted to the ORU Titan cluster as a single-node job. Thus, *bt* jobs are submitted to Titan when the whole-genome calculation is performed.

Once all the pairwise comparisons have been performed, a second program, *MCount*, is run to merge the results for each chromosome’s match output binary vector for all of its comparisons to the other genome’s chromosomes. For example, for a given base chromosome, *B*, the *p* comparisons for that chromosome are combined, such that for each position in the binary vector, if a 1 is found in any of the *p* binary vectors for the base chromosome, then a match for that position is counted. The output is the total number of matches and the total number of base pairs for the base chromosome. Considering, for instance, these two binary vectors, 0011000101 and 1000101010, the merged vector in this case would be 1011101111, the count would be 8, and the size would be 10. A minimum match size is specified for each analysis.

GeneCompare utilizes an indexing of the base chromosome, recording the positions of every unique 16-bp prefix found in the chromosome. Then, for each 16-bp prefix in the target chromosome that it finds, it starts a comparison at each location that prefix was found in the base chromosome, starting with a match size of 16 and then, from there, continuing base-pair-by-base-pair comparisons to determine if a match of the minimum size *m* is found. All matches found result in 1’s being recorded in the binary vectors for both the base and target chromosomes. The code is multi-threaded via OpenMP with the searches being fully parallel, taking advantage of the many cores on each of Titan’s nodes. To validate the efficacy of the algorithm, individual chromosomes were compared to themselves (i.e., *H. sapiens* chromosome 1 to *H. sapiens* chromosome 1). These tests yielded a 100% match.

### Nucleotide BLAST

Further validations for sequence comparisons generated by the GeneCompare program utilized the National Center for Biotechnology Information (NCBI) Basic Local Alignment Search Tool (BLAST) (Johnson et al., 2008) (BLAST: Basic Local Alignment Search Tool (nih.gov)), specifically nucleotide BLAST. Nucleotide BLAST was applied to compare the generated sequence locations from *P. paniscus* and *H. sapiens*. The BLAST alignment entailed aligning pairs of sequences, with each letter in each sequence matched from base pair to base pair and gaps introduced in non-matching regions (Bergman, 2007). To perform this task, the respective *H. sapiens* Hg14.38 and *P. paniscus* Mhudiblu_PPA_v0 reference genomes were selected. The choice of the chromosome reference number depended on the chromosome where matching sequences were found in both *P. paniscus* and *H. sapiens*. The start location and length of each matched sequence were determined using the GeneCompare results. To calculate the end location, the length of the match was added to each start location. The *P. paniscus* and *H. sapiens* sequences alternated as the subject and query in the comparisons, with the analysis run twice for each case. During the BLAST analysis, the standard databases were chosen, and the program was optimized for highly similar sequences (mega BLAST). Algorithm parameters were maintained in their default settings, including a maximum of 100 target sequences, automatic adjustment of short queries for short input sequences, an expected threshold of 0.05, a word size of 28, and a maximum number of matches in the query range of 0. Scoring parameters encompassed match/mismatch scores of 1 and -2, along with linear gap costs. Filters and masks were applied, specifically selecting options for low-complexity regions and masking for the lookup table only. For each BLAST run, an alignment score was generated by assigning values to each aligned pair and subsequently averaging these individual scores over the entire alignment. Based on the alignment score and visual representation of sequence alignments, the results of the GeneCompare algorithm could be further validated. Once the comparisons were validated, the obtained data were meticulously recorded and annotated, considering available gene annotations.

### WashU Epigenome Browser

The validation process of motif sequences discovered within the human genome involved the utilization of the Washington University in St. Louis (WashU) Epigenome Browser (WashU Epigenome Browser (wustl.edu)). Typically employed for the interactive exploration of genomic data through a web browser interface (Li et al., 2022), this browser was directed towards the Hg14.38 genome. Upon initiation, the JASPAR Transcription Factors 2022 collection was integrated. This involved selecting “annotation tracks” within the larger section of “tracks.” Once this was established, the JASPAR Transcription Factors were added from the displayed drop-down menu. In adding the transcription factors, the generated binding site locations were measured against known binding sites. Other methods utilized in verifying the motif sequences used were chromatin immunoprecipitation sequencing (ChIP-seq) and RNA sequencing (RNA-seq). This entailed using ChIP-seq and RNA-seq tracks from available public data hubs on the WashU browser. This consisted of tracks from mammary epithelial cells and peripheral blood mononuclear cells, among many others. The strategic inclusion of transcription factors facilitated a comparative analysis of select motif sequences against established binding sites. The validation extended to motifs exhibiting consistent recurrence within proximate locales, prompting additional rounds of validation to determine precision and reliability.

## Results

Utilizing an alignment-dependent algorithm, matches of 32 base pairs or greater were queried. Validation was performed using BLAST for specific sequences to verify the legitimacy of the GeneCompare results. *H. sapiens* to *H. sapiens* genomes were also run against each other using GeneCompare to verify high similarity within a species. The dependent variable, the *H. sapiens* reference genome, was the base against which the *P. paniscus* genome, the independent variable, was tested. The corresponding chromosomes of each species tested against each other are shown in Table 1. “Matched Pairs” represent the numerical amount of total base pairs matched between the two chromosomes, “Total Pairs” is the numerical length of base pairs in the *P. paniscus* chromosome, and “Percent Ratio” is the ratio between Matched Pairs and Total Pairs, expressed as a percentage.

**Table 1.** Alignment-dependent *Homo sapiens* against *Pan paniscus* chromosome match results at 32-bp granularity. Each chromosome matched between the two species, the total number of matched characters, and the overall percentage similarity.

| *H. sapiens*<br>Chromosome<br>Number | *H. sapiens*<br>Chromosome<br>Length | *P. paniscus*<br>Chromosome<br>Number | Matched<br>Pairs | Total Pairs | Percent<br>Ratio |
| --- | --- | --- | --- | --- | --- |
| 1 | 248,956,422 | 1 | 198830515 | 224621958 | 88.52% |
| 2 | 242,193,529 | 2A | 86476631 | 104947789 | 82.40% |
| 2 | - | 2B | 96032718 | 129381034 | 74.22% |
| 3 | 198,295,559 | 3 | 179859032 | 195577393 | 91.96% |
| 4 | 190,214,555 | 4 | 137405367 | 182437434 | 75.32% |
| 5 | 181,538,259 | 5 | 105303732 | 176502593 | 59.66% |
| 6 | 170,805,979 | 6 | 153998744 | 168932342 | 91.16% |
| 7 | 159,345,973 | 7 | 128407208 | 150536359 | 85.30% |
| 8 | 145,138,636 | 8 | 120262480 | 141842281 | 84.79% |
| 9 | 138,394,717 | 9 | 85693932 | 109767803 | 78.07% |
| 10 | 133,797,422 | 10 | 115087820 | 128853861 | 89.32% |
| 11 | 135,086,622 | 11 | 118227986 | 129867894 | 91.04% |
| 12 | 133,275,309 | 12 | 86385936 | 131319602 | 65.78% |
| 13 | 114,364,328 | 13 | 86164223 | 95736914 | 90.00% |
| 14 | 107,043,718 | 14 | 80220476 | 87894197 | 91.27% |
| 15 | 101,991,189 | 15 | 66105775 | 80799215 | 81.81% |
| 16 | 90,338,345 | 16 | 52717439 | 71000456 | 74.25% |
| 17 | 83,257,441 | 17 | 47011016 | 77747126 | 60.47% |
| 18 | 80,373,285 | 18 | 56247446 | 74093087 | 75.91% |
| 19 | 58,617,616 | 19 | 48805932 | 55604062 | 87.77% |
| 20 | 64,444,167 | 20 | 53861088 | 59769695 | 90.11% |
| 21 | 46,709,983 | 21 | 29685737 | 33144400 | 89.56% |
| 22 | 50,818,468 | 22 | 27913273 | 31064846 | 89.85% |
| X | 156,040,895 | X | 134107583 | 146233785 | 91.71% |

**Table 2.** Alignment-dependent *Homo sapiens* against *Pan paniscus* chromosome match results at 200-bp granularity. Each chromosome matched between the two species, the total number of matched characters, and the overall percentage similarity.

| *H. sapiens*<br>Chromosome<br>Number | *H. sapiens*<br>Chromosome<br>Length | *P. paniscus*<br>Chromosome<br>Number | Matched<br>Pairs | Total Pairs | Percent<br>Ratio |
| --- | --- | --- | --- | --- | --- |
| 1 | 248,956,422 | 1 | 65924683 | 248956422 | 26.48% |
| 2 | 242,193,529 | 2A | 25367055 | 242193529 | 10.47% |
| 2 | - | 2B | 30309715 | 242193529 | 12.51% |
| 3 | 198,295,559 | 3 | 56304045 | 198295559 | 28.39% |
| 4 | 190,214,555 | 4 | 38927642 | 190214555 | 20.47% |
| 5 | 181,538,259 | 5 | 28359116 | 181538259 | 15.62% |
| 6 | 170,805,979 | 6 | 48352397 | 170805979 | 28.31% |
| 7 | 159,345,973 | 7 | 37715657 | 159345973 | 23.67% |
| 8 | 145,138,636 | 8 | 35724351 | 145138636 | 24.61% |
| 9 | 138,394,717 | 9 | 24357045 | 138394717 | 17.60% |
| 10 | 133,797,422 | 10 | 35451654 | 133797422 | 26.50% |
| 11 | 135,086,622 | 11 | 36958574 | 135086622 | 27.36% |
| 12 | 133,275,309 | 12 | 24763551 | 133275309 | 18.58% |
| 13 | 114,364,328 | 13 | 25659765 | 114364328 | 22.44% |
| 14 | 107,043,718 | 14 | 25446479 | 107043718 | 23.77% |
| 15 | 101,991,189 | 15 | 20232985 | 101991189 | 19.84% |
| 16 | 90,338,345 | 16 | 12619516 | 90338345 | 13.97% |
| 17 | 83,257,441 | 17 | 12354429 | 83257441 | 14.84% |
| 18 | 80,373,285 | 18 | 16489418 | 80373285 | 20.52% |
| 19 | 58,617,616 | 19 | 12627026 | 58617616 | 21.54% |
| 20 | 64,444,167 | 20 | 16022239 | 64444167 | 24.86% |
| 21 | 46,709,983 | 21 | 7937341 | 46709983 | 16.99% |
| 22 | 50,818,468 | 22 | 7874479 | 50818468 | 15.50% |

**Table 3.** Alignment-dependent *Homo sapiens* against *Pan paniscus* chromosome match results at 1000-bp granularity. Each chromosome matched between the two species, the total number of matched characters, and the overall percentage similarity.

| *H. sapiens*<br>Chromosome<br>Number | *H. sapiens*<br>Chromosome<br>Length | *P. paniscus*<br>Chromosome<br>Number | Matched<br>Pairs | Total Pairs | Percent<br>Ratio |
| --- | --- | --- | --- | --- | --- |
| 1 | 248,956,422 | 1 | 214723 | 248956422 | 0.09% |
| 2 | 242,193,529 | 2A | 94195 | 242193529 | 0.04% |
| 2 | - | 2B | 59666 | 242193529 | 0.02% |
| 3 | 198,295,559 | 3 | 139968 | 198295559 | 0.07% |
| 4 | 190,214,555 | 4 | 74690 | 190214555 | 0.04% |
| 5 | 181,538,259 | 5 | 65315 | 181538259 | 0.04% |
| 6 | 170,805,979 | 6 | 128356 | 170805979 | 0.08% |
| 7 | 159,345,973 | 7 | 94621 | 159345973 | 0.06% |
| 8 | 145,138,636 | 8 | 83180 | 145138636 | 0.06% |
| 9 | 138,394,717 | 9 | 54092 | 138394717 | 0.04% |
| 10 | 133,797,422 | 10 | 91954 | 133797422 | 0.07% |
| 11 | 135,086,622 | 11 | 95700 | 135086622 | 0.07% |
| 12 | 133,275,309 | 12 | 53388 | 133275309 | 0.04% |
| 13 | 114,364,328 | 13 | 51472 | 114364328 | 0.05% |
| 14 | 107,043,718 | 14 | 89421 | 107043718 | 0.08% |
| 15 | 101,991,189 | 15 | 40239 | 101991189 | 0.04% |
| 16 | 90,338,345 | 16 | 23942 | 90338345 | 0.03% |
| 17 | 83,257,441 | 17 | 30128 | 83257441 | 0.04% |
| 18 | 80,373,285 | 18 | 42890 | 80373285 | 0.05% |
| 19 | 58,617,616 | 19 | 57217 | 58617616 | 0.10% |
| 20 | 64,444,167 | 20 | 37723 | 64444167 | 0.06% |
| 21 | 46,709,983 | 21 | 11140 | 46709983 | 0.02% |
| 22 | 50,818,468 | 22 | 22414 | 50818468 | 0.04% |
| X | 156,040,895 | X | 388126 | 156040895 | 0.25% |

Using GeneCompare provided a comprehensive view of genome comparisons among higher primate species. Several search lengths were used, including a 32-bp, 200-bp, and 1000-bp algorithm. All possible chromosome matches were tried. However, consistently the expected chromosomes were found to be the highest matches, i.e., chromosome 1 matched with chromosome 1. With the 32-bp search, chromosome 3 had the highest similarity between the two species at 91.96%, and chromosome 5 had the lowest match at 59.66% similarity (Table 1). As expected, the lowest granularity tested, 32 base pairs, yielded the highest percentage matches between the two species. However, switching from 32-bp granularity to 200-bp granularity revealed interesting divergences in percent similarity. For instance, chromosome 16 is a 74.25% match at the 32-bp granularity, but drops to a 13.97% match at 200 bp, effectively diminishing it from a moderate match to one of the lowest chromosome matches.

Contrastingly, chromosomes 1, 6, and 11 show the highest percentage similarities at both the 32-bp granularity and at the 200-bp granularity. These results indicate that chromosome-to-chromosome comparisons prove more indicative of relatedness than averaged genome-to-genome comparisons. Furthermore, the original molecular studies on protein similarity and these results highlight the need to explore regulatory sequences of DNA and their role in species development.

Preliminary pattern analysis of DNA motifs for transcription factor binding sites such as KLF4, TBP, and SRY indicate that these consensus sequences are conserved across the two species. These transcription factor binding sites were chosen due to their universal regulatory roles within the genome. Kruppel-like factor 4 (KLF4) is known to regulate cell growth, proliferation, and differentiation in pluripotent stem cells (Ghaleb & Yang, 2017). TATA-box binding protein (TBP) is known as a universal transcription factor involved in forming the transcription initiation complex (Davidson et al., 2004; Hantsche & Cramer, 2017). Sex-determining region Y protein (SRY) is a transcription factor that has been implicated in the epithelial-mesenchymal transition, as well as male sex determination during early stages of development (Wilhelm et al., 2007; Wu et al., 2020). The preliminary data suggest that the number of appearances of the KLF4 binding sites between the two species is 97.12% identical, in line with the original protein studies. Likewise, TBP and SRY appearances are 89.51% and 98.9% identical, respectively. Further studies into highly repetitive motifs within higher primates, and even within the same species will shed light onto the role repetitive sequences play in gene expression. While it is valuable to explore the similarity of protein-coding DNA sequences, and even whole genome sequences between species, understanding how genomic information is regulated is also important, complemented with whole transcriptome and proteome studies.

## Discussion

The results obtained using GeneCompare challenge the widely cited figure of 98.4% genomic similarity between humans and bonobos (Sibley et al., 1990), revealing a substantially lower percentage of shared sequences when considering the entire genome. This discrepancy likely stems from the GeneCompare algorithm’s stringency, its inclusion of non-protein-coding regions, and its lack of preselection for highly conserved gene families. The approach presented in this paper provides a holistic view of genomic similarity, revealing considerable variation in homology across different chromosomes independently of biases towards chromosome length or protein-coding sequences.

The observation that certain chromosomes, such as chromosome 3, exhibit significantly higher similarity than others, such as chromosome 5, highlights the importance of analyzing chromosome-specific homology rather than relying on averaged genome-wide comparisons. This heterogeneity suggests that different genomic regions have experienced varying rates of evolution and may be subject to different selective pressures. Further investigation is warranted to understand the underlying mechanisms driving these differences. Potential factors could include varying rates of mutation, recombination, gene duplication, transposition, and horizontal gene transfer.

The analysis of sequence matches at different word sizes (32-bp vs. 200-bp) further emphasizes the complexity of genomic comparisons. The reduction in similarity observed for chromosome 16 when shifting from 32-bp to 200-bp granularity suggests that this chromosome contains a significant number of short, interspersed homologous regions rather than extended stretches of conserved sequence. This type of analysis can shed light on the types of evolutionary changes that have occurred and their impact on overall genomic similarity. Additionally, the preliminary analysis of transcription factor binding sites (KLF4, TBP, and SRY) reveals a high degree of conservation in these regulatory regions between humans and bonobos. The nearly identical occurrence of these binding sites is consistent with the previously reported high protein sequence similarity between the two species (King & Wilson, 1975). This finding supports the known functional importance of these regulatory elements and their role in maintaining essential cellular processes. Future studies should investigate the context of these binding sites within the genome, the specific transcription factors expressed, and the resulting gene expression patterns to fully understand the functional implications of this conservation.

While GeneCompare provides a valuable tool for identifying homologous sequences, its linear search-based approach may be computationally intensive for very large genomes and could be further optimized for speed. The validation using NCBI BLAST, while helpful, is limited by the inherent biases of alignment-based methods. Future work should focus on expanding the phylogenetic scope of this study to include other higher primate species such as *G. gorilla, Pongo orangutan*, and *Pan troglodytes*. Such comparisons would provide a more complete picture of genomic divergence within the Hominidae family. Although higher primates share notable similarities in their anatomical and physiological structures, as well as in their proteomes, their whole genomes can differ more than these phenotypic characteristics would suggest (Mao et al., 2021; Prüfer et al., 2012). Ultimately, a multi-faceted approach that combines genomic, transcriptomic, proteomic, and epigenetic data will be essential for unraveling the complex history of humans and their closest relatives.

## Acknowledgments

This work was supported by NSF Grant 2201435.

## Author contributions

C.R., D.D., and E.A. carried out all of the genome analysis experiments. R.N. and S.W. developed sequencing algorithms and maintained the TITAN supercomputer. M.G. assisted in figure design and manuscript editing. W.R. provided the original research idea. J.G. conceived the project, interpreted the results, and wrote the manuscript. E.A., E.D., Y.A., and C.V. assisted in manuscript writing and editing.

## Data and software availability statement

The datasets generated and programs in the current study are available from the corresponding authors for reasonable requests.

---

**Figure 1.**
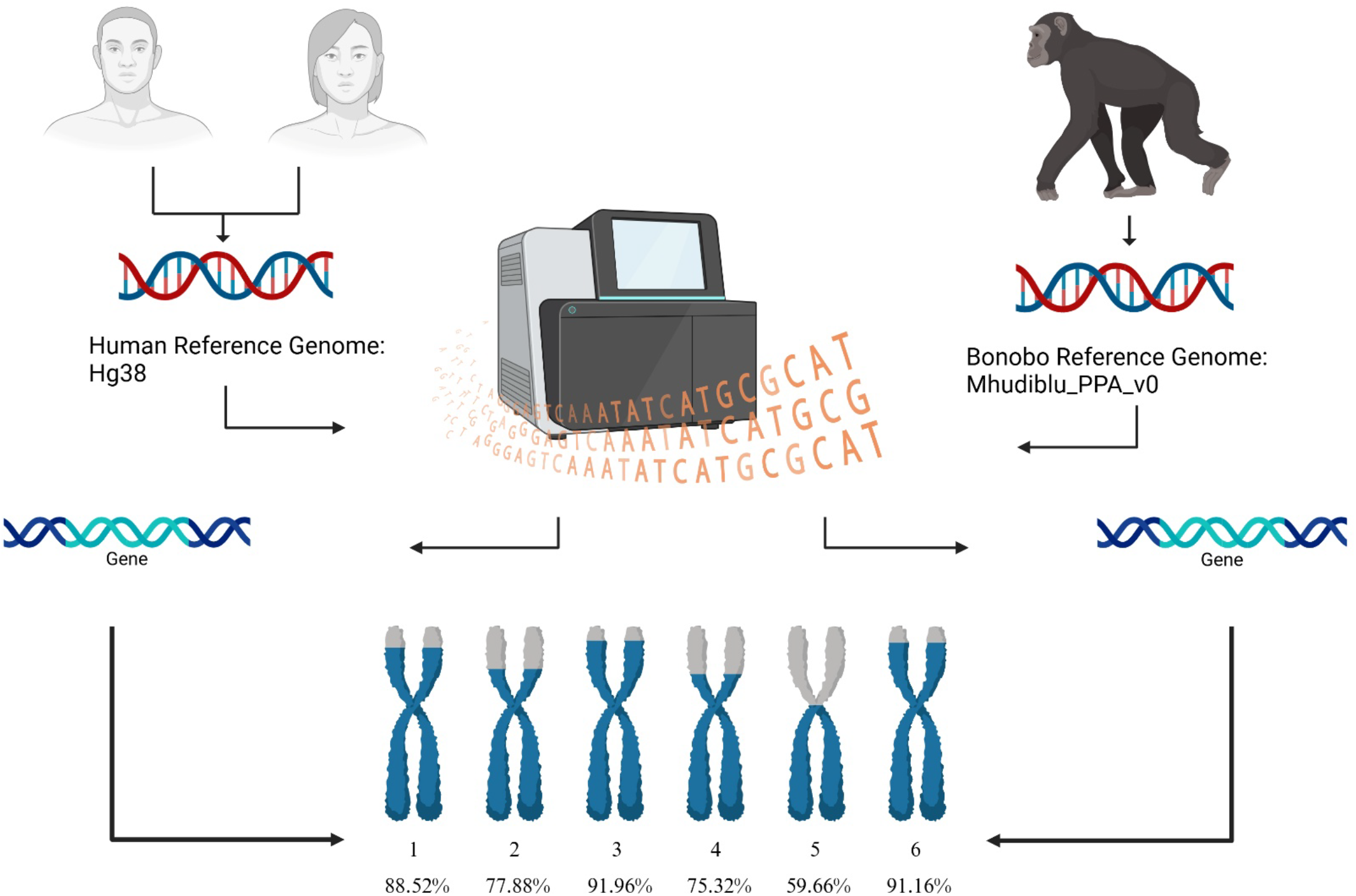
Genome comparison between *Pan paniscus* (Mhudiblu_PPA_v0) and *Homo sapiens* (Hg38.p13) reference genomes. Depiction of a comprehensive whole genome comparison between the *P. paniscus* and *H. sapiens* genomes, utilizing the human reference genome Hg38.13 and bonobo reference genome Mhudiblu_PPA_v0. The comparison identified similarity matches in both coding and noncoding regions of the DNA. The percentages that are indicated under each chromosome represent the proportion of matches found between the two organisms. (Created with Biorender.com)

**Figure 2.**
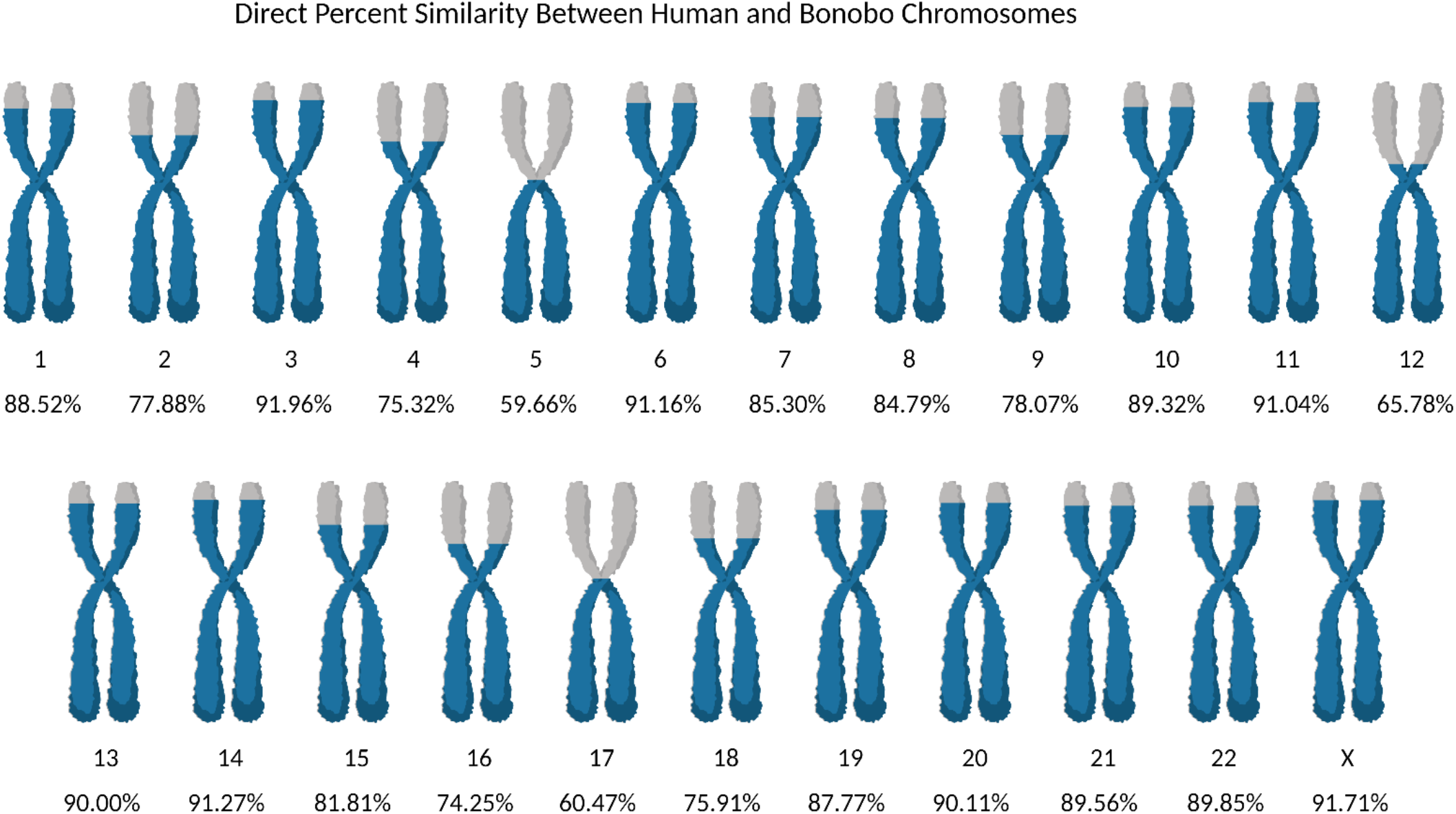
DNA percentage match between *Homo sapiens* against *Pan paniscus* chromosomes. Individual chromosomes between *H. sapiens* and *P. paniscus* genomes were aligned using a 32bp matching algorithm. Overall, the chromosomes ranged from 91.96% similar to 59.66% similar. Chromosome 3 was the most similar and chromosome 5 was the most dissimilar between the two species. Created with Biorender.com

**Figure 3.**
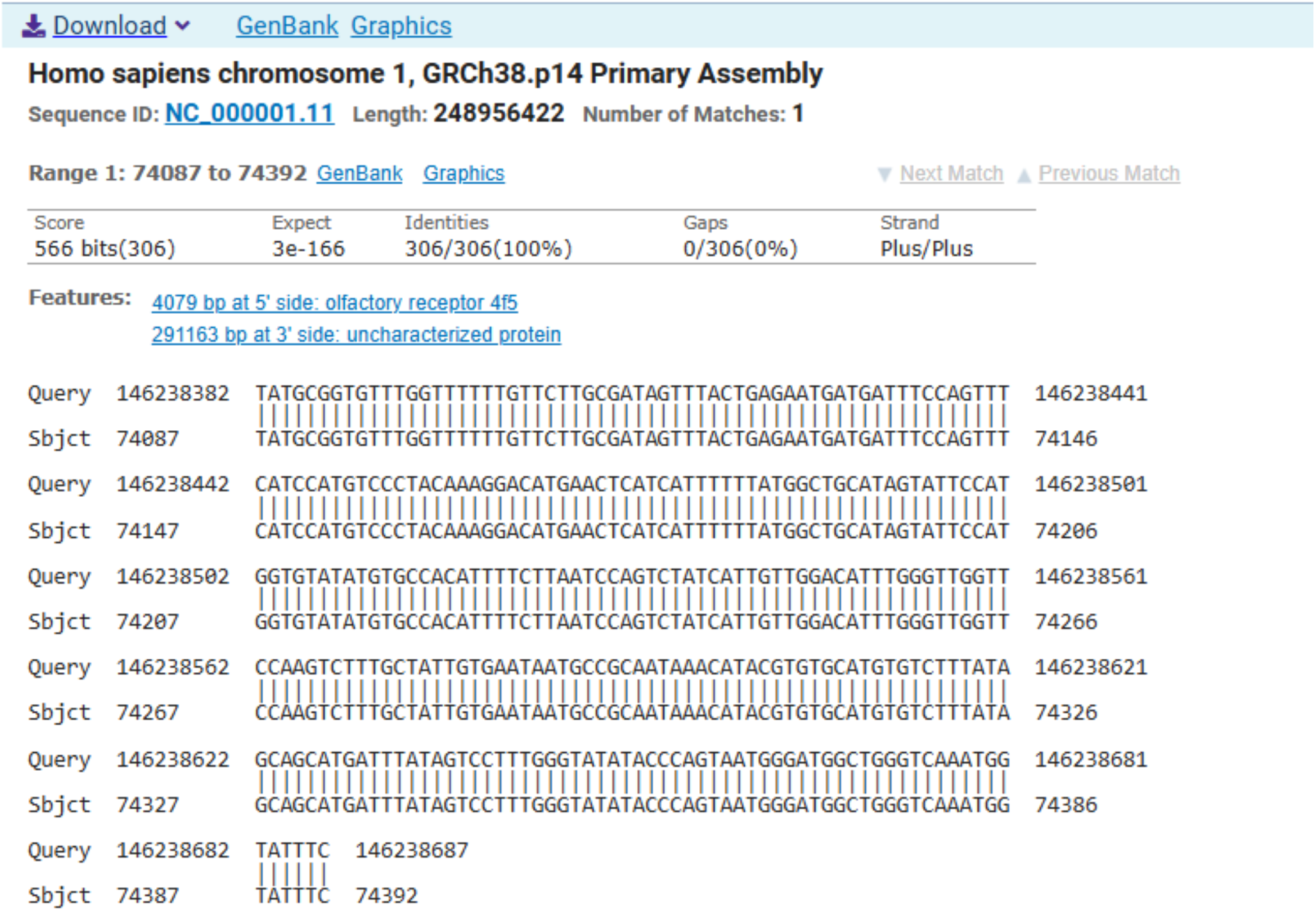
NCBI BLAST validation of sequence comparisons. The figure above depicts the outcome of the NCBI BLAST analysis performed on genomic sequences obtained from *P. paniscus* (Bonobo) and *H. sapiens* (Human) chromosome 1. In this analysis, the query sequence consisted of *P. paniscu*s loci spanning positions 146,238,382 to 146,238,688, while the subject sequence comprised *H. sapiens* loci ranging from positions 74,087 to 74,393. The results of the analysis revealed a 100% sequence identity between the corresponding loci, which is visually depicted by connecting lines indicating matched base pairs. Furthermore, notable findings indicate the presence of a match within the *H. sapiens* protein-coding gene known as olfactory receptor 4F5 (OR4F5) in the 5’ direction and an uncharacterized protein-coding gene in the 3’ direction.

